# Distributed and multifaceted effects of threat and safety

**DOI:** 10.1101/2021.07.11.451944

**Authors:** Kelly Morrow, Murty Dinavahi, Jongwan Kim, Song-tao Song, Kesong Hu, Luiz Pessoa

## Abstract

Sustained anticipation of unpredictable aversive events generates anticipatory processing that is central to anxiety. In the present functional Magnetic Resonance Study (fMRI) study, we examined how sustained threat is processed in the human brain. We used a relatively large sample (*N* = 109) and employed a Bayesian multilevel analysis approach to contrast `threat` and `safe` periods. Our analyses demonstrated that the effect of sustained threat is heterogeneous and distributed across the brain. Thus, the impact of threat is widespread, and not restricted to a small set of putatively emotion-related regions, such as the amygdala and the bed nucleus of the stria terminalis. Both transient and sustained, and increased and decreased responses during threat were observed. Our study reveals that transitioning between threat and safe states, and vice versa, leads to a widespread switch in brain responding that involves most of the brain.

---

Anxiety/fear related mental health conditions afflict a large number of individuals. In humans, the neural basis of threat processing has been intensively investigated in the last two decades. Anxious states induced by acute, unpredictable stressors have widespread effects on brain function due to changes in the organization of large-scale brain networks (Hermans et al., 2011, 2014; Kienast et al., 2008; Scott et al., 2006; Thomason et al., 2011)

Despite progress, several important questions remain unanswered:

1. If threat is <u>sustained</u> over periods of time (say, over 10 seconds), do brain responses exhibit sustained responses over the time interval in question? In other words, it is important to determine if a region’s response is *transient* or *sustained* (Alvarez et al., 2011; Hasler et al., 2007; Hur et al., 2020; McMenamin et al., 2014; Schlund et al., 2013; Somerville et al., 2013).
2. Nonhuman research indicates that the <u>amygdala</u> plays a role in processing transient “fear”-related stimuli (e.g., a condiotioned stimulus) but the <u>bed nucleus of the stria terminalis</u> (BST) is implicated in sustained “anxious” processing (Davis and Shi, 1999; Walker et al., 2003). Such putative division of labor between the two structures is now debated (Canteras et al., 2009; Klumpers et al., 2017; Paré and Quirk, 2017; Tovote et al., 2015; Shackman and Fox, 2016).
3. Threat-related processing engages multiple brain regions, such that responses are stronger during threat compared to neutral conditions. However, growing evidence reveals that a complementary set of regions respond <u>less strongly</u> during threat relative to safe. For example, in a recent study, regions along the midline (including the posterior cingulate cortex, PCC) showed decreased responses for proximal vs. distal threat (Meyer et al., 2019), consistent with previous findings of the virtual tarantula paradigm by (Mobbs et al., 2010); see also (Yao et al., 2018). In particular, many regions that exhibit threat-related signal decreases appear to overlap with the default network. Thus, it is important to establish the relationship between threat-related decreases and the regions of the default network.
4. Most studies of temporally extended threat have performed analyses <u>assuming that hemodynamic responses follow a canonical shape</u>. Whereas this assumption is often reasonable for experimental paradigms with brief events, it is problematic when longer conditions are employed that generate responses that do not follow those based on the canonical shape; see, e.g., (Chen et al., 2015; Sreenivasan and D’Esposito, 2019). The ability to estimate response shape is particularly important when characterizing whether or not threat-related responses are transient or not (Hur et al., 2020).
5. Many previous studies of extended threat processing involved relatively modest sample sizes, and focused their analysis at the voxel level, a combination that results in relatively <u>limited statistical power</u> (Button et al., 2013; Cremers et al., 2017; Turner et al., 2018).
6. Another gap in the literature that is likely related to limited statistical power is <u>the focus of studies on a relatively small set of brain regions</u>, even though studies are performed at the whole-brain level. Because of this, we still lack important knowledge about the involvement of specific brain regions in the human brain during extended threat processing. For example, in rodents, the ventral striatum has been implicated in some forms of threat processing (Boeke et al., 2017), and sectors of the hippocampus play important roles in threat-related processing (Canteras et al., 2009; Tovote et al., 2015). As a further example, now in humans, although studies have reported threat-related responses in the cerebellum (Lange et al., 2015; Moreno-Rius, 2018), this structure has not been investigated in a more targeted manner.

To address these gaps in the literature, we studied *N* = 109 participants during an uncertain-threat paradigm (Figure 1). We focused on a set of 85 brain regions that have been discussed in the literature in the context of threat-related processing (Figure 2), and estimated the responses during `threat` and `safe` blocks without making response-shape assumptions.

**Figure 1:**
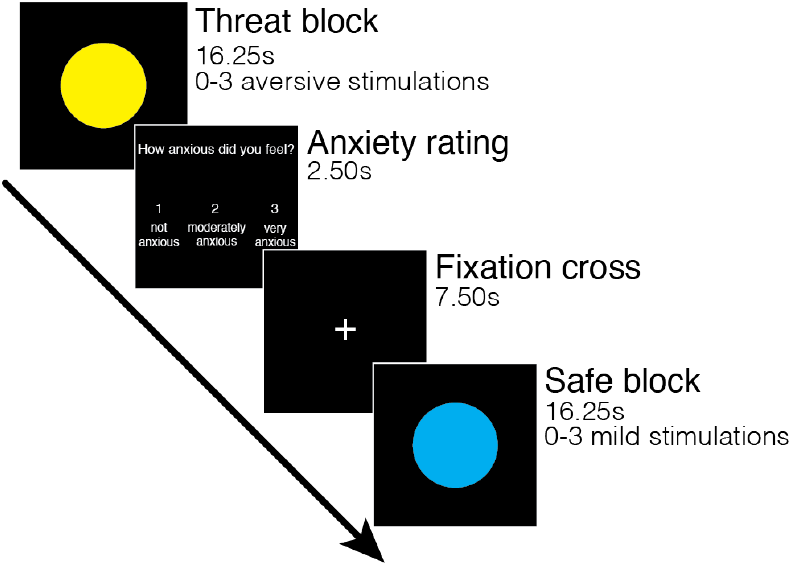
Paradigm. During the `threat` condition, signaled by a yellow circle, participants could receive 0-3 aversive stimulations (shocks) over a period of 16.25 seconds. During the `safe` condition, signaled by a blue circle, participants could receive 0-3 mild stimulations (vibration). After each block, participants rated their anxiety levels on a scale of 1-3, followed by a 7.50 second fixation cross before the next block.

**Figure 2:**
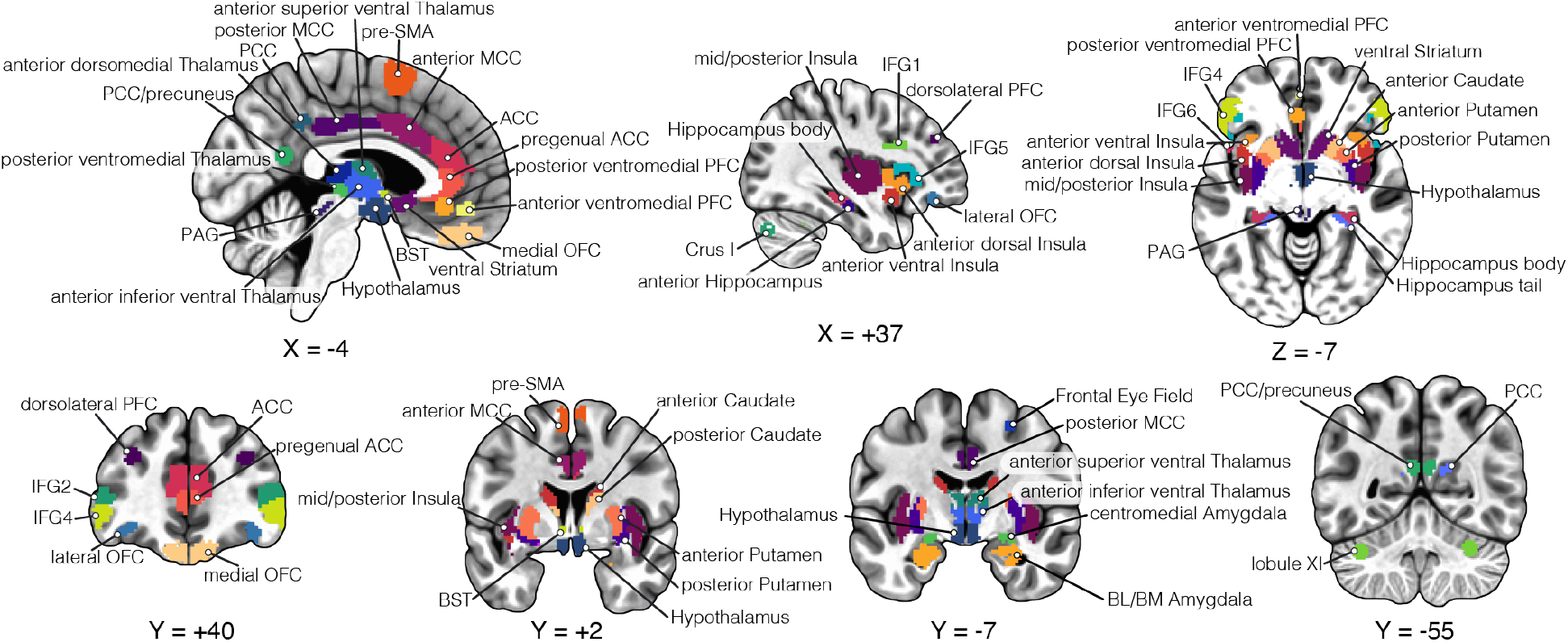
Regions of interest. Brain regions analyzed. Abbreviations: ACC; anterior cingulate cortex, MCC; midcingulate cortex, PCC; posterior cingulate cortex, BL/BM amygdala; basolateral/basomedial amygdala, BST; bed nucleus of the stria terminalis, IFG; inferior frontal gyrus, OFC; orbitofrontal cortex, PAG; periaqueductal gray, PFC; prefrontal cortex, pre-SMA; pre-supplementary motor area.

We would like to highlight our statistical approach; see also (Chen et al., 2019; Gelman et al., 2012; Gelman and Hill, 2006; Limbachia et al., 2021). Whole-brain analysis with fMRI often lacks statistical power to uncover effects at the voxel level, which can lead to poor replicability. Therefore, here we sought to focus on a set of carefully delineated regions of interest (ROIs) and leverage the strengths of Bayesian multilevel modeling (Gelman and Hill, 2006; McElreath, 2018) (BML). One of the strengths of BML is that it allows the simultaneous estimation of multiple clustered parameters within a single model (in an educational setting, for example, the effects at multiple schools within a district). In the present context, BML allowed the estimation of the effects across multiple ROIs simultaneously, or voxels simultaneously (Chen et al., 2019). Among the advantages of this approach, information about the effect in one region/voxel can be shared across all regions/voxels (technically referred to as “partial pooling”).

## Methods

### Participants

One hundred and nine right-handed participants (53 females; ages 18-35 years; average: 21.17 years, standard deviation: 2.59 years) with normal hearing, normal or corrected-to-normal vision, and no reported neurological disease or current us of psychoactive drug use were recruited from the University of Maryland, College Park, community. The study was approved by the University of Maryland, College Park, Institutional Review Board and all participants provided written informed consent before participating in the study. All were paid immediately after the experiment.

### Stimuli and behavioral paradigm

To induce a sustained anxious states, a threat of shock paradigm was used while participants were scanned. Prior to scanning, participants completed the Trait Anxiety portion of the Spielberger State-Trait Anxiety Inventory (STAI) (Spielberger et al., 1970), and completed the State Anxiety portion immediately before scanning.

The experiment consisted of 3 runs over a two-hour session. Each run consisted of 8 `threat` and 8 `safe` blocks presented randomly. On each block, a colored circle was presented for 16.25 seconds via PsychoPy (www.psychopy.org). Prior to scanning, participants were informed that the color of the circle indicated whether a block was `threat` (yellow circle) or `safe` (blue circle). A `threat` circle indicated that they could receive zero to three highly unpleasant, but not painful electrical stimulations (referred to as “shock” here), whereas a `safe` circle indicated that they could receive zero to three very mild, but perceptible stimulations (referred to as “touch”).

Both stimulation levels were set by the participant to reflect the desired intensities prior to scanning, and adjusted between runs, if needed. After each 16.25-second block, participants rated how anxious they felt during the block on a scale of 1-3 (1: not anxious, 2: moderately anxious, and 3: very anxious). Anxiety ratings were followed by a 7.50 second fixation cross before proceeding to the next trial. Because the ratings used by participants were largely correlated with block condition (i.e., higher ratings were indicated during `threat` blocks), we did not analyze ratings here.

Of the 48 trials in total, 32 trials (16 `threat` and 16 `safe`) contained zero stimulations and the remaining 16 trials (8 `threat` and 8 `safe`) had at least one instance of stimulation.

### MRI data acquisition

MRI data collection used a 3T Siemens TRIO scanner (Siemens Medical Systems) with a 32-channel head coil. First, we acquired a high-resolution T1-weighted MPRAGE anatomical scan (TR: 2400 ms, TE: 2.01 ms, FOV: 256 mm, voxel size: 0.8-mm isotropic). Subsequently, we collected functional echo planar image (EPI) volumes over three runs (340 images per run) using a multiband scanning sequence with TR = 1250 ms, TE = 39.4 ms, FOV = 210 mm, and multiband factor = 6. Each volume contained 66 non-overlapping oblique slices oriented 30 degrees clockwise relative to the AC-PC axis; thus voxels were 2.2 mm isotropic. Double-echo field maps (TE = 73.0 ms) were also acquired with acquisition parameters matched to the functional data.

### Skin conductance response acquisition

Skin conductance responses (SCR) were collected using the MP-150 data acquisition system (BIOPAC Systems, Inc., Goleta, CA) with the GSR100C module. Signals were acquired at 250 Hz using MRI-compatible electrodes attached to the index and middle fingers of the participants’ non-dominant, left hand. SCR data from two subjects (1 female, 1 male) were not collected due to technical problems.

### Regions of interest (ROIs)

We focused on 85 structurally and functionally defined cortical and subcortical regions of interest (shown in Figures 3 and 4). Functional masks were based on data from separate studies, not the present one. Potential contributions of white matter and cerebrospinal fluid were minimized by excluding voxels that intersected with masks of these tissue types. All ROIs were disjoint, such that no overlap occurred between them.

**Figure 3:**
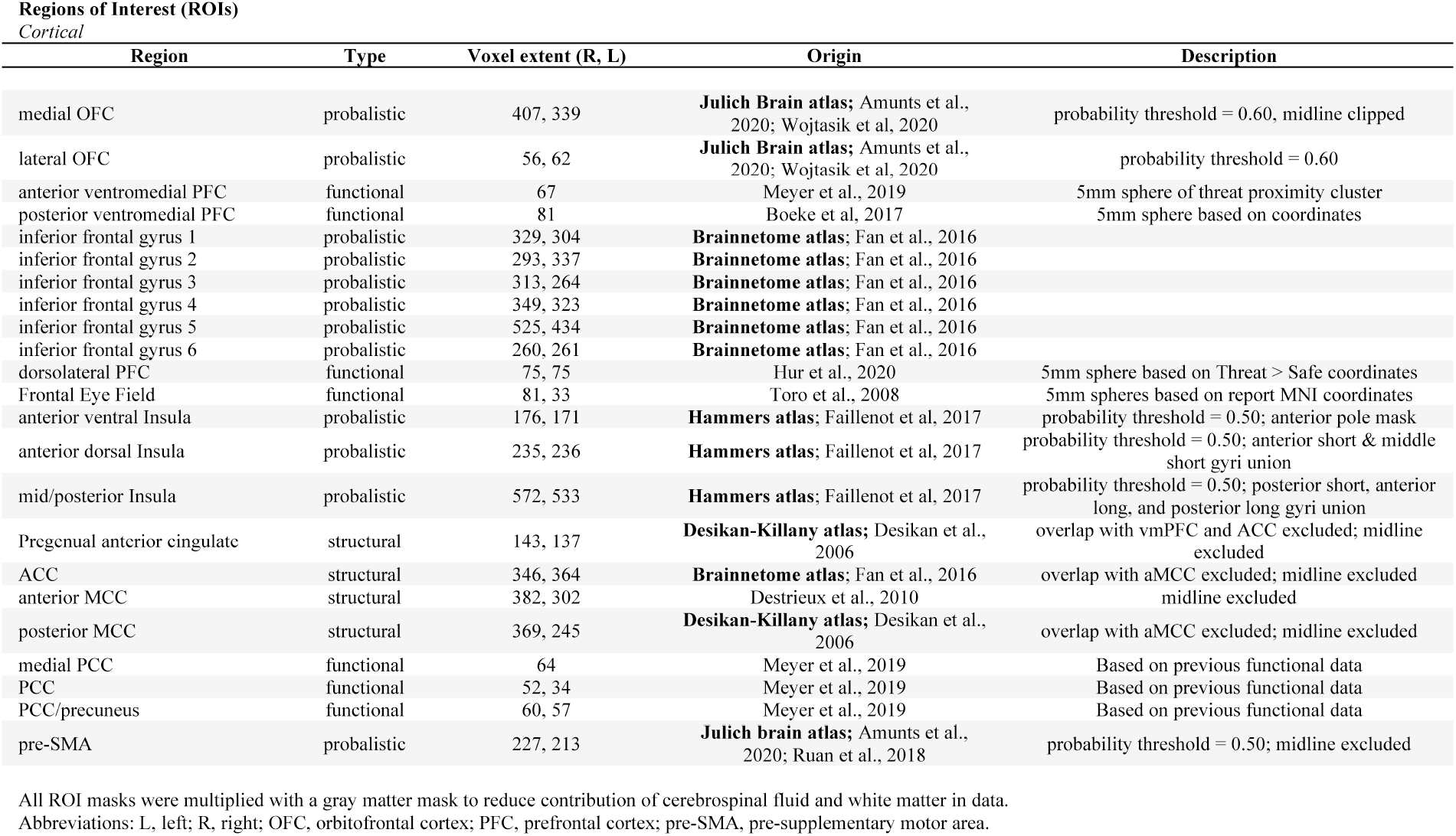
Cortical regions of interest.

**Figure 4:**
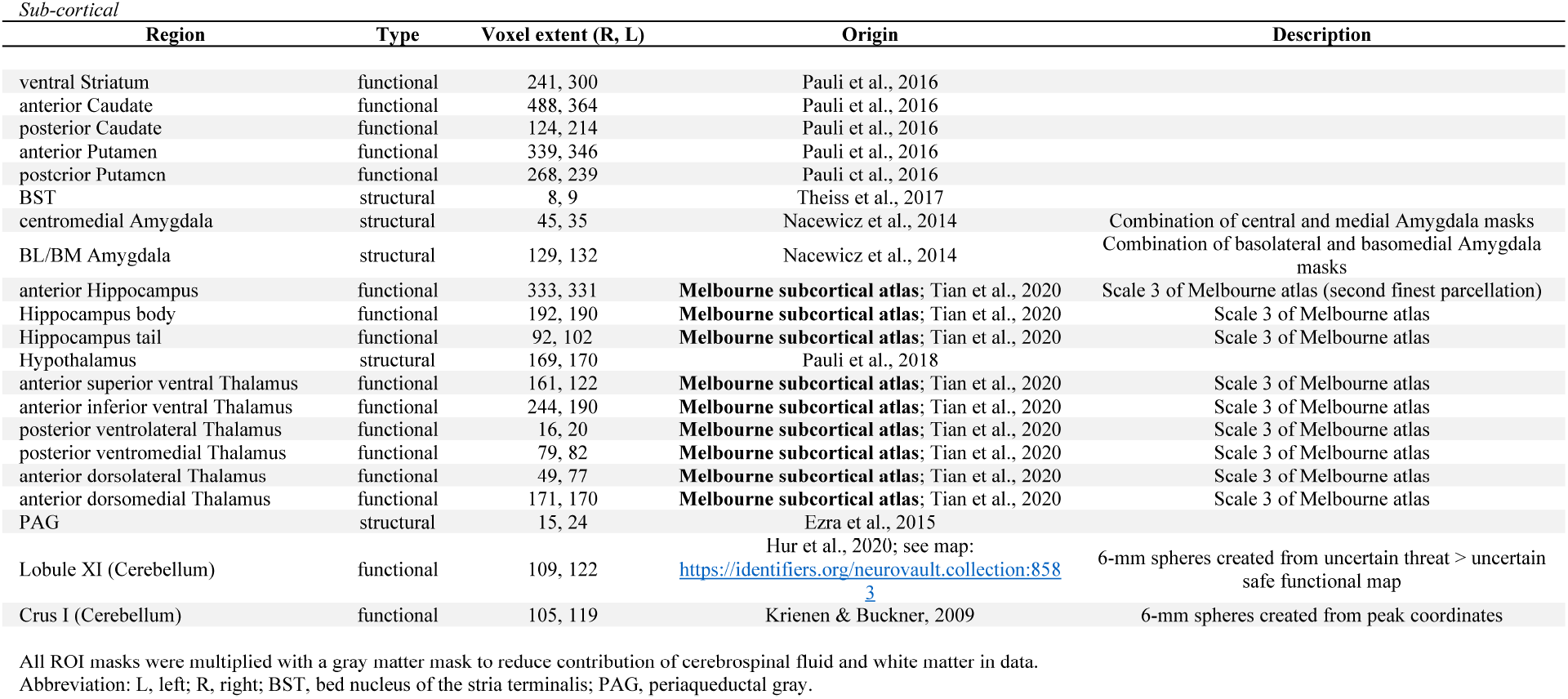
Subcortical regions of interest.

### Data preprocessing

Functional images were preprocessed as described in our previous work by using a combination of fMRI packages and in-house scripts (Limbachia et al., 2021). Slice-timing correction was performed with the Analysis of Functional Neuroimages (AFNI; (Cox, 1996)) `3dTshift` with Fourier interpolation to align the onset times of each slice in a volume to the first acquired slice. For voxelwise analysis (but not ROI level), data were spatially smoothed with a Gaussian filter (4 mm full-width half-maximum) restricted to gray matter voxels. Signal intensity at each voxel was normalized to a mean value of 100 separately for each run. To further reduce the contribution of head motion artifacts, FSL’s Independent Component Analysis, Automatic Removal of Motion Artifacts (ICA-AROMA) toolbox was applied (Pruim et al., 2015). Components classified as head motion were regressed out of the data using FSL’s `fsl_regfilt`.

Given the small size of some of our regions of interest, following previous work (Limbachia et al., 2021), we sought to improve coregistration between functional and anatomical images using a “voting scheme” to determine whether or not a voxel belonged to the brain (i.e., skull stripping). To do so, we employed six different fMRI packages (ANTs: Avants et al. (2009), AFNI: Cox (1996), ROBEX: Iglesias et al. (2011), FSL: Smith et al. (2004), SPM: Friston et al. (2007), and BrainSuite: Shattuck and Leahy (2002)). Based on T1 structural data, if 4/6 packages estimated a voxel to belong to the brain, it was retained, otherwise it was discarded. Next, ANTs was used to estimate a nonlinear transformation mapping the skull-stripped anatomical T1 image to the skull-stripped MNI152 template (interpolated to 1-mm isotropic voxels). The nonlinear transformations from coregistration/unwarping and normalization were combined into a single transformation that was applied to map volume-registered functional volumes to standard space (interpolated to 2-mm isotropic voxels).

Preprocessed functional data were then subjected to run-wise motion thresholding. Any particular run that exceeded a mean frame-wise displacement of 0.5 mm, had a total framewise displacement of 5 mm or more, or had 20% or more of all TRs with framewise displacement over 0.5 mm was excluded. Runs were then concatenated for each subject.

Temporal signal-to-noise ratio, defined as the ratio of mean to standard deviation across time, can be rather low in some brain areas, including the orbitofrontal cortex, amygdala, and periaqueductal gray. Voxels that did not meet our criteria were discarded from analysis (0.21 ± 0.47% of voxels). Voxels were excluded if their mean (after signal normalization to 100) was outside the range 95-105, or the standard deviation exceeded 25.

### Subject-level analysis

To estimate the shape of the responses during `threat` and `safe` blocks, we focused on the 16 blocks of each block type with no electrical stimulation. Response shape was estimated by employing a series of cubic spline basis functions (similar to the “finite impulse response” method also commonly adopted). The analysis utilized subject-level multiple linear regression with AFNI’s `3dREMLfit` program.

At the ROI level, the analysis employed the average timeseries (unsmoothed data) across ROI voxels. At the voxel level, spatially smoothed data were used to increase signal to noise, as routinely done in fMRI analysis. Blocks of a given type were modeled with 14 regressors (i.e., cubic splines) aligned to the TRs of the block starting at block onset (so starting at *t* = 0 and ending with the TR starting at *t* = 16.25).

The anxiety rating period, which was not of interest here, was modeled by convolving a square wave with a canonical hemodynamic response. Likewise, the remaining blocks containing at least one stimulation event were modeled by convolving the block duration (16.25 seconds) with a canonical filter. To minimize contamination of task-related signal with electrical stimulation events, the 15 seconds following each stimulation were censored from analysis.

Separately, we estimated the responses to stimulation events to confirm the presence of robust responses in regions like the amygdala. In this case, the cubic spline regressors were aligned to the stimulation (shock or touch) onset. Finally, all analyses included a set of regressors modeling head motion parameters (six rigid body motion parameters and their discrete temporal derivatives), in addition to linear and nonlinear polynomial terms (up to third order) to account for baseline and slow signal drift.

### Group Bayesian multilevel analysis: ROI level

Employing a <u>single multilevel</u> model allowed the contributions to fMRI signals of subject-level effects (i.e., subject effect across conditions) and ROI-level effects (i.e., ROI effect across subjects) to be estimated simultaneously with the condition effect (`threat` versus `safe`). The input to the model consisted of 85 averaged-across-voxels time series from the first level described above, as in our previous work (Limbachia et al., 2021). The model was estimated using the `brms` R package (Bürkner, 2017) which utilizes `rstan` (https://mc-stan.org/users/interfaces/rstan), the R interface to the `Stan` proba-bilistic language (https://mc-stan.org).

We considered two temporal time windows: `early` (2.5-8.75 seconds from block onset) and `late` (10-16.25 seconds). Note that responses at 0 and 1.25 seconds were not used to account for the latency of the hemodynamic response. For each window, we defined <u>response magnitude</u> as the sum of the responses across the window. In doing so, our goal was to create something akin to an “area under the curve” response measure (Chen et al., 2015). However, experimental conditions do not have a common baseline across subjects or ROIs. Thus, response estimates at each time point (per subject and ROI) will potentially have negative and positive values across time that may “cancel out” each other if summed across the temporal window of interest, potentially leading to response magnitudes close to zero because negative and positive values were simply summed (see, for example, Figure 10). Accordingly, to determine response measures, response estimates were minimum-shifted, resulting in positive values only, for each subject and ROI, separately (note that the original estimated responses are shown in the figures).

Given our within-subject design, we used the <u>difference</u> between `threat` and `safe` response magnitudes as the variable to be tested: Δ_*s*,*r*_, where *s* indexes subjects, and *r* indexes ROI. For computational expediency, we analyzed the two time windows separately taking into account anxiety-related covariates. In standard linear mixed-effects modeling notation our model was defined as

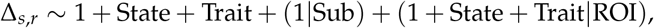

where Sub (Subject) and ROI are grouping variables. State and Trait indicate the covariates of the model. The terms “1” indicate intercept terms, including an overall intercept and so-called varying intercepts per subject (subject-specific contribution) and per ROI (ROI-specific contribution). The notation (1 + State + Trait|ROI) summarizes the varying intercepts and slopes per ROI, but also the correlation structure between intercept and slopes.

Model estimation generates a <u>single posterior</u> that is a joint distribution in a high-dimensional parameter space. Posteriors for each effect at the ROI-level were calculated by summing the relevant contributions (i.e., overall intercept term and intercept at the ROI level). Note whereas posteriors can be plotted separately for each ROI for visualization purposes (Figures 7 and 8), they should be understood as belonging to a single joint distribution.

Bayesian estimation requires the specification of prior distributions for all the parameters of interest in the model. We employed so-called weakly informative priors that help estimation convergence (all 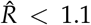 < 1.1, as recommended), but have negligible impact on estimated parameters, especially with reasonable sample sizes. We provide a complete description of the Bayesian formulation of the model, together with the priors, in the following GitHub page: https://github.com/LCE-UMD/mood-anxiety.

### Group Bayesian multilevel analysis: Voxel level

We performed voxel-level BML analysis of insula and amygdala data, separately. The analysis entailed adding a voxel level to the model above. In addition, instead of using the original ROI definitions, we created finer parcellations for the insula and amygdala. Thus, the 941 voxels of the entire left insula were clustered into 11 ROIs, and the 986 voxels of entire right insula were clustered into 10 ROIs. Similarly, the 167 voxels of the entire left amygdala were clustered into 6 ROIs, and the 176 voxels of entire right amygdala were clustered into 6 ROIs. In all cases, the new smaller ROIs respected the original ROI boundaries investigated in the region-level analysis (e.g., new insula ROIs did not include voxels from both the ventral and dorsal insula; or correspondingly, from the centromedial and the basolateral amygdala). Smaller ROIs were created by clustering adjacent voxels based on their *{x*, *y*, *z}* coordinates (not time series data) with standard *k*-means clustering.

The motivation for having finer parcellations of the insula and amygdala was related to the concept of partial pooling in multilevel modeling. By having a larger number of ROIs, the pooling effect (that is, voxel effects possibly being pushed to some extent to the overall average of the ROI), was more strongly restricted to the local ROI to which a voxel belonged, thus respecting the idea that information exchange should tend to stay local (technically speaking, partial pooling depends on the variance of voxel-level estimates). Overall, the modeling approach followed the same strategy as that of the the ROI-level analysis, with the addition of the voxel-level data:

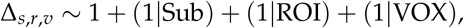

where *s*, *r*, and *v* index subject, (small) ROI, and voxel, respectively. For computational expediency, covariates were not included in the models. In addition, for the insula, separate models were applied to the left and right insula. Because the number of voxels in the amygdala was relatively small, we included an additional hemisphere level in the equation above, and a single model was run. Again, convergence of estimates was observed in all cases (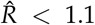 < 1.1, as recommended).

### Skin conductance

As stated, data were acquired at 250 Hz. For each run, we detrended the signal by removing the best linear fit to the data. Subsequently, we applied a median filter over 50-sample windows (200 ms) to remove high-frequency noise.

## Results

We report the results of Bayesian multilevel analysis (BML) performed on 85 regions (one representative time series per ROI obtained by averaging unsmoothed functional data to preserve spatial resolution). Currently, whole-brain voxelwise BML analysis is not computationally feasible. Accordingly, we performed voxelwise analysis in two regions important for threat-related processing, namely the insula and the amygdala. In all Bayesian analyses, statistical evidence for effects is reported in terms of *P*+, the probability that the effect is greater than zero based on the posterior distribution: values closer to 1 provide evidence that the effect of interest is greater than zero (`threat` > `safe`) while values closer to zero convey support for the inverse effect (`safe` > `threat`). We treat Bayesian probability values as providing a continuous amount of support for a given hypothesis; thus not dichotomously as “significant” vs. “not significant”. ^1^

Because Bayesian multilevel modeling implements so-called <u>partial pooling</u> of the estimates across voxels (when present), regions, as well as participants, the process tends to generate estimates that are more conservative; for example, they will be closer to the average effect within a given region than if each voxel’s effect were estimated individually. Effectively, the approach allows information to be shared across spatial units and tends to stabilize the effect estimates. Because of this conservative nature, some statisticians have suggested that adjustment for multiplicity is not needed (Gelman et al., 2012), especially since all the inferences are drawn from a single, overall posterior distribution of an integrative model. We follow this approach here; for detailed discussion, see (Chen et al., 2021).

### Hemodynamic responses to shock and skin conductance responses during threat blocks

To verify that shock delivery evoked clear hemodynamic responses, we estimated responses evoked by stimulus administration. Large, transient responses were observed across many brain regions (Figure 5; note the scale of the responses). Note that robust differential responses were not observed in the basal amygdala.

**Figure 5:**
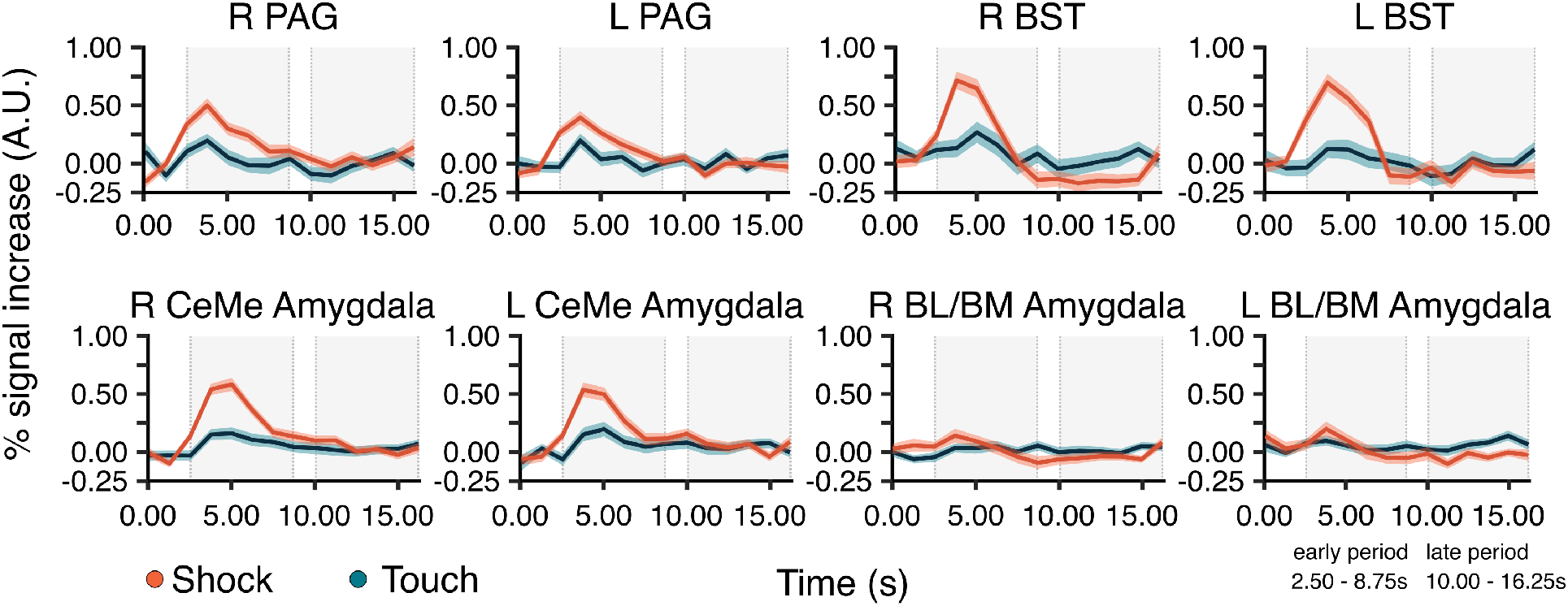
Hemodynamic responses to shock and touch. Estimated responses averaged across subjects. Error bands show 95% interval for standard error across participants to illustrate variability only. Shaded regions represent early and late periods. Abbreviations: L: Left, R: Right, PAG: periaqueductal grey, BST: bed nucleus of stria terminalis, CM: centromedial, BL/BM: basolateral/basomedial.

Analysis of skin conductance revealed transient responses following the onset of `threat` blocks that was not seen in `safe` blocks (Figure 6). It is noteworthy that, although stimulation events spanned the duration of a block, autonomic arousal indexed by skin conductance response was not sustained.

**Figure 6:**
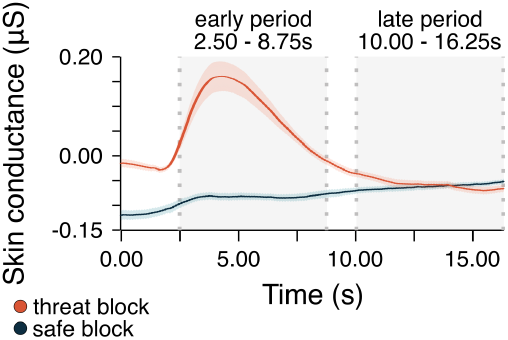
Skin conductance responses for safe and threat blocks. Skin conductance responses averaged across 107 subjects (solid lines) for safe (dark blue) and threat (red) blocks. Error bands show 95% interval for standard error across participants to illustrate variability only. Shaded regions represent early and late periods.

**Figure 7:**
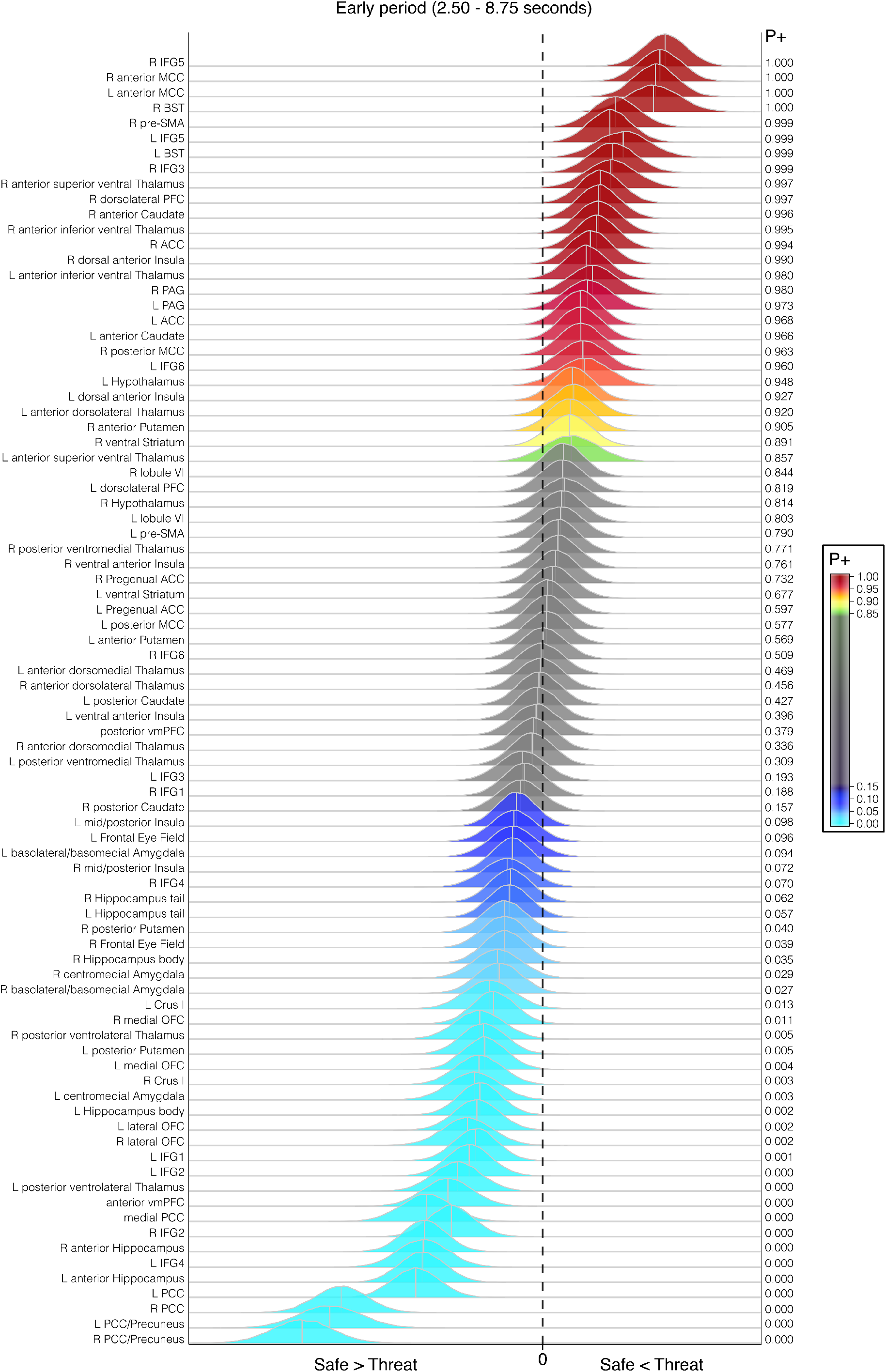
Posterior distributions: Early period. The posteriors show the distribution of the difference between `threat` and `safe`. *P*+ is the probability that the effect is greater than zero.

**Figure 8:**
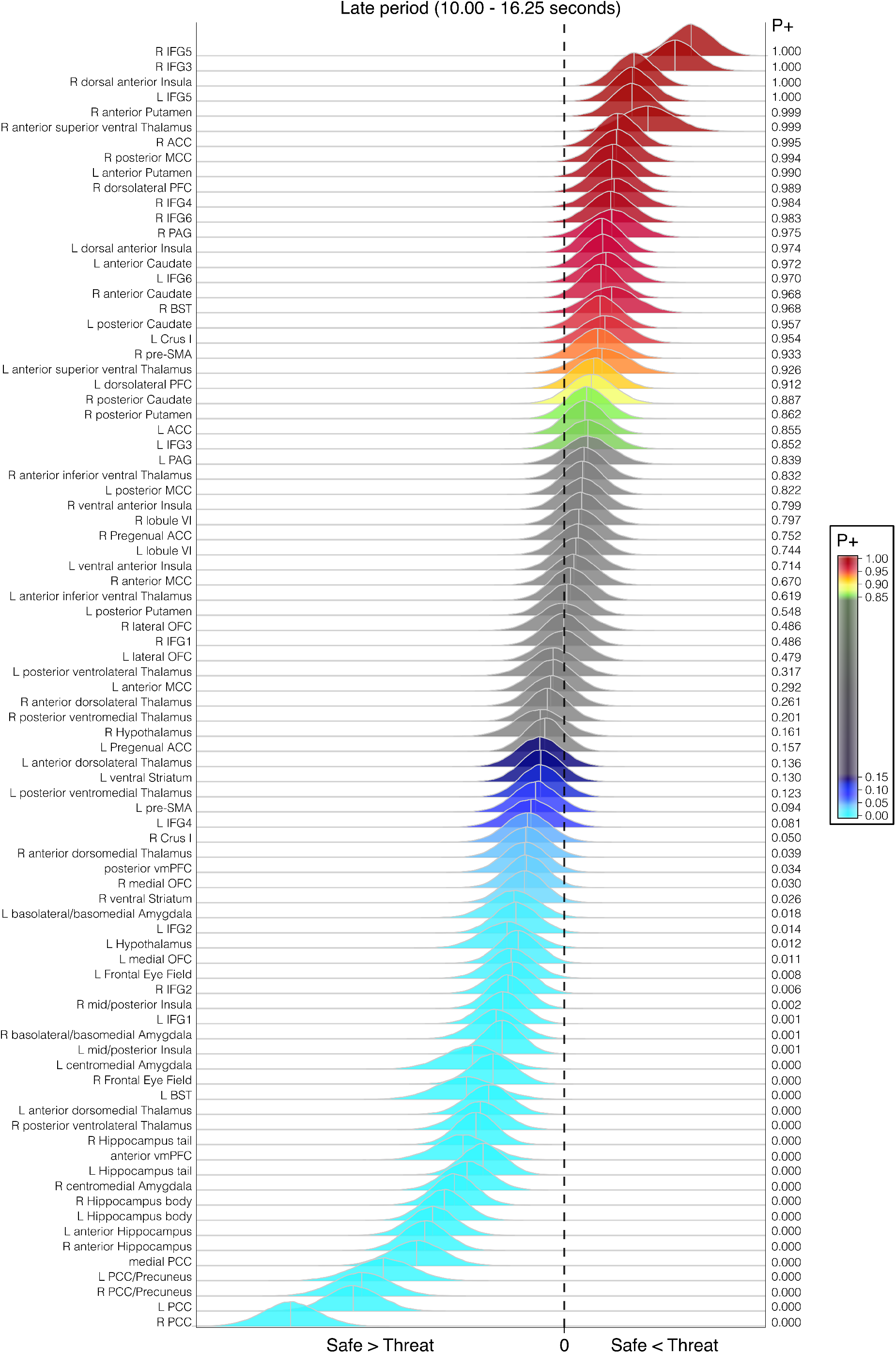
Posterior distributions: Late period. The posteriors show the distribution of the difference between `threat` and `safe`. *P*+ is the probability that the effect is greater than zero.

### ROI-level analysis

A broad set of brain regions involved in multiple aspects of threat-related processing were considered for analyses (Figure 2). For each ROI, responses were characterized based on two temporal windows of a block: an `early` window from 2.50 to 8.75 seconds and a `late` window from 10.00 to 16.25 seconds after block onset. The average response from each window was considered the response strength for a given condition.

The results are summarized via a series of posterior distributions, one for each brain region (Figures 7 and 8). To illustrate these results, Figure 9 shows probability values on brain slices for the `early` and `late` periods. We see very strong support (say, *P*+ > 0.990) for `threat` > `safe` in both early and late periods for multiple regions, as well as strong statistical support for many other regions. A similar picture was observed for `threat` < `safe` effects (this time with *P*+ values very close to zero).

**Figure 9:**
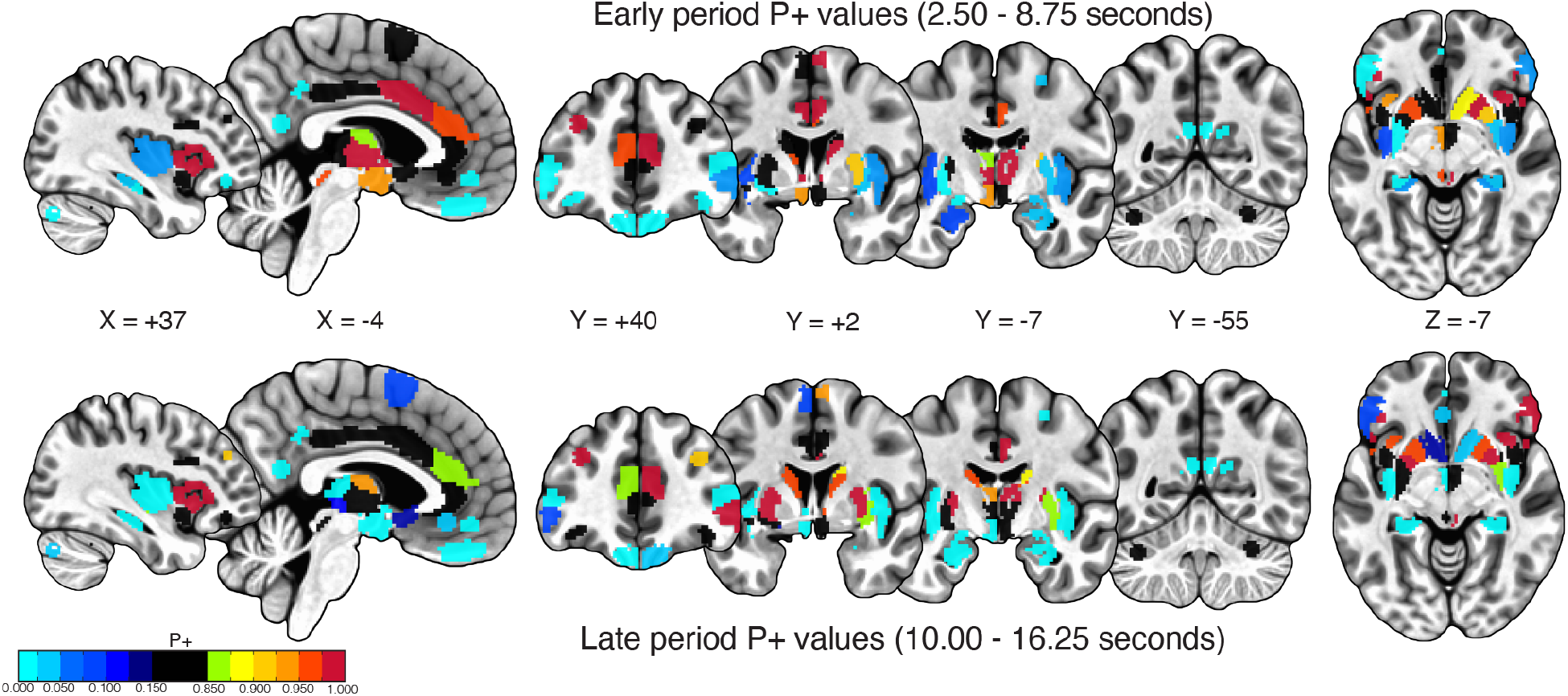
Contrast of threat vs. safe. Probability that `threat` > `safe` effect is great than zero during early and late periods. Values closer to 1 indicate greater responses during threat blocks, and values closer to 0 indicate greater activity during safe blocks. Brain slices correspond to those in Figure 2.

**Figure 10:**
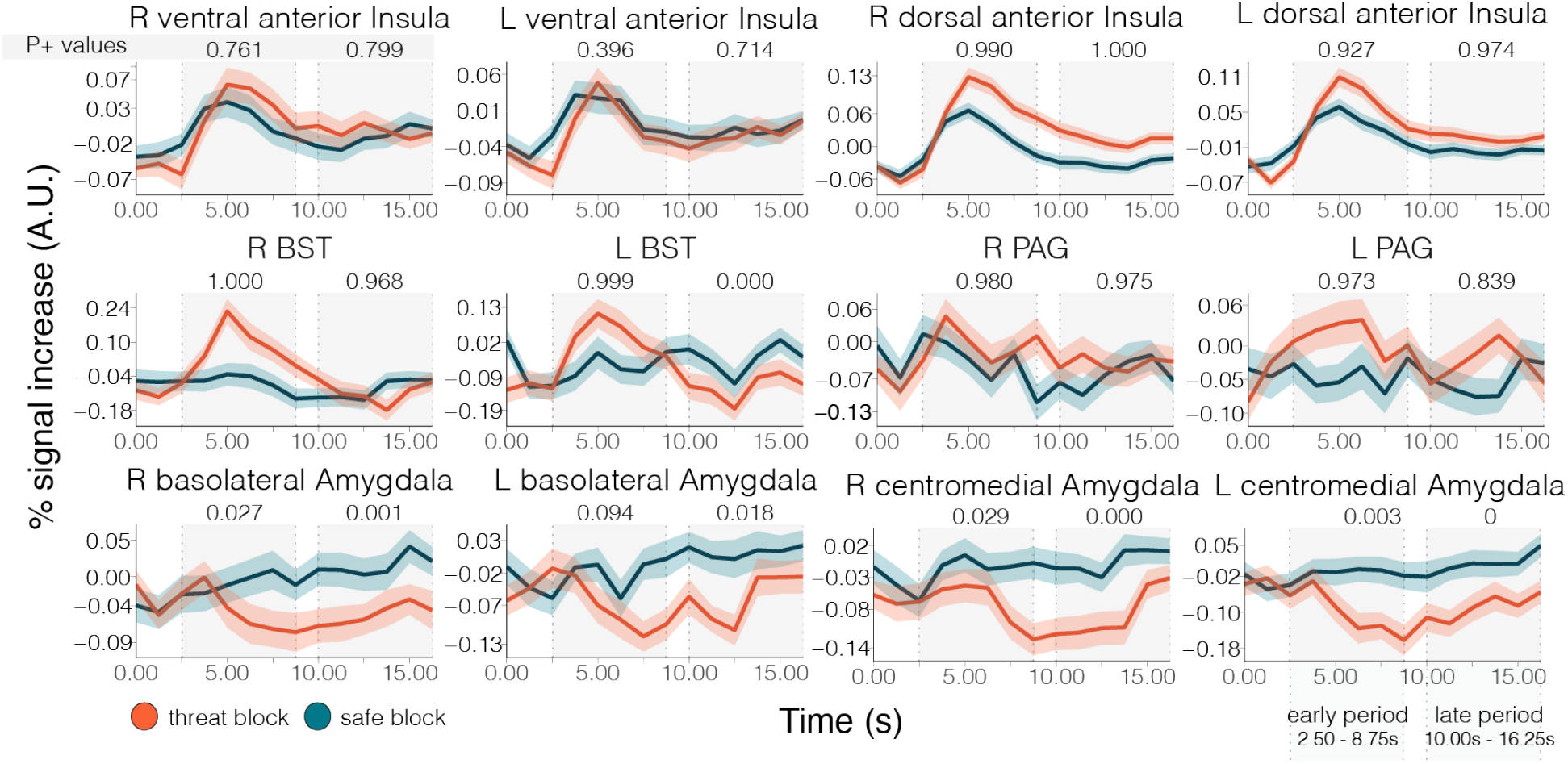
Estimated responses across key regions. Error bands show 95% interval for standard error across participants to illustrate variability only.

We show estimated responses across multiple sets of regions to illustrate the types of responses observed, including transient and sustained responses. Figure 10 shows responses in a few key brain regions. We observed robust response decreases during threat, which we illustrate in Figure 11. An additional set of “noteworthy” regions is displayed in Figure 12. Figure 13 shows responses in the striatum. Finally, Figure 14 shows responses in the thalamus.

**Figure 11:**
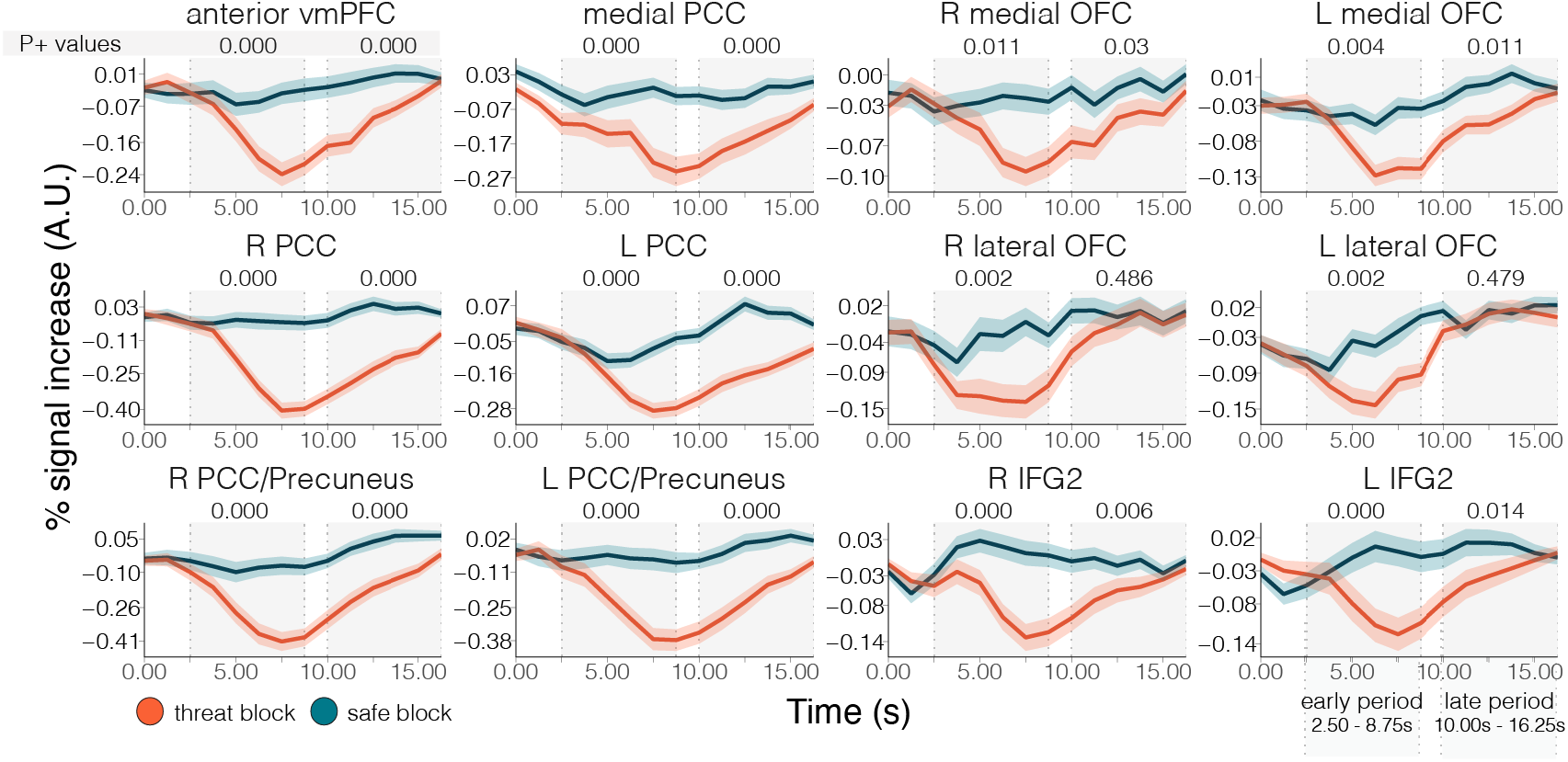
Estimated responses across regions exhibiting decreased responses during threat. Error bands show 95% interval for standard error across participants to illustrate variability only.

**Figure 12:**
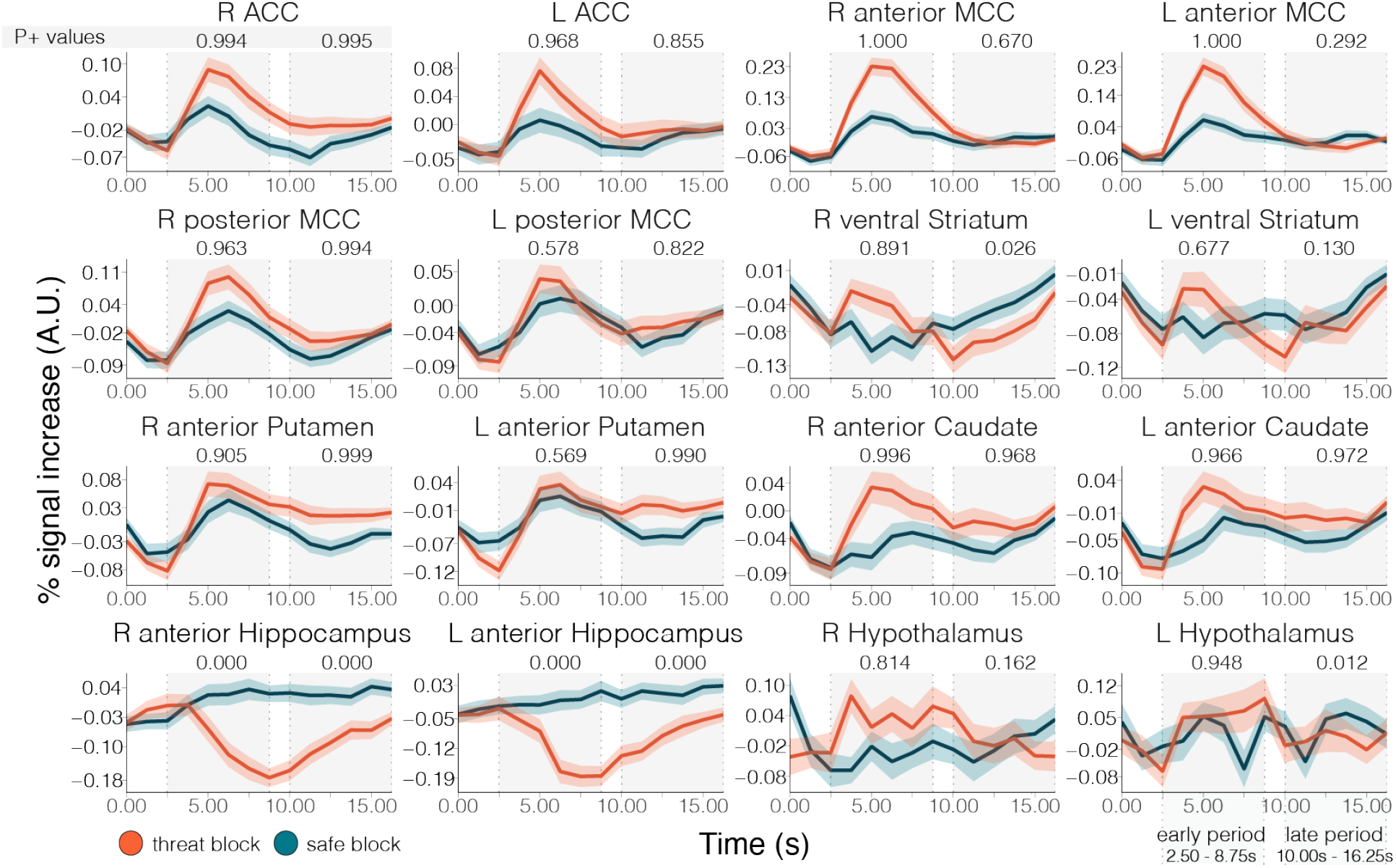
Estimated responses across additional regions. Error bands show 95% interval for standard error across participants to illustrate variability only.

**Figure 13:**
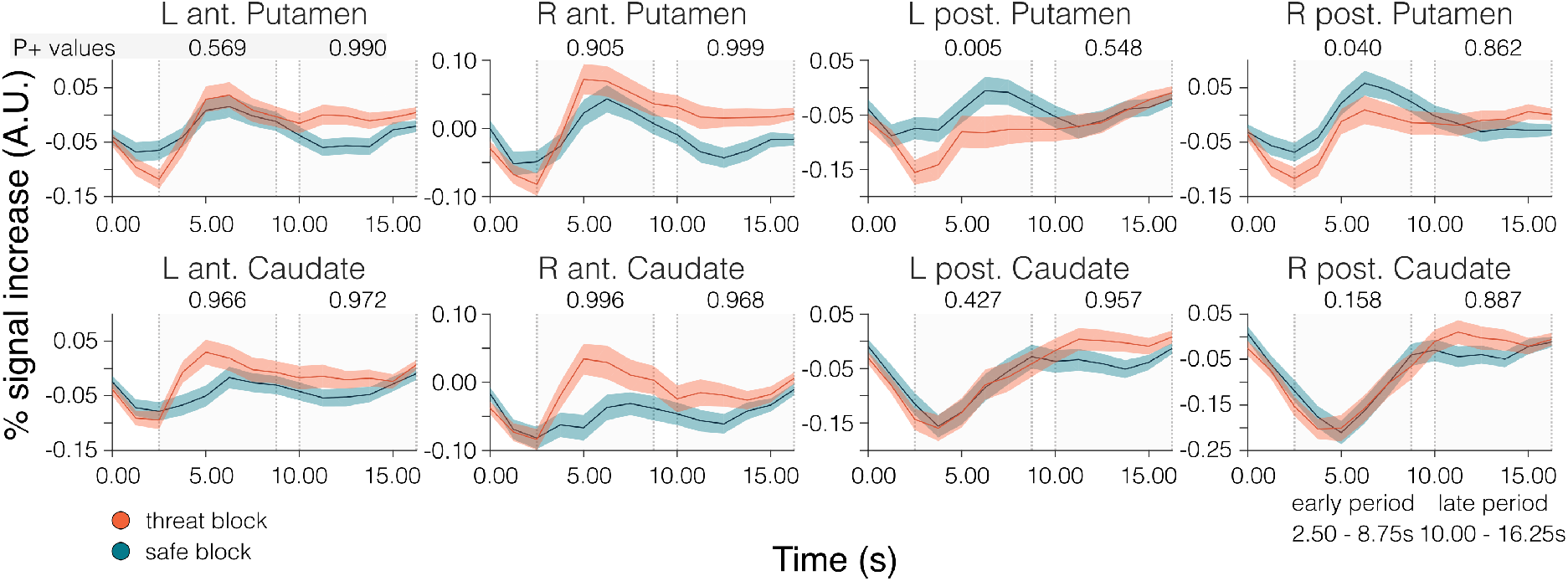
Estimated responses for cau-date and putamen. Error bands show 95% interval for standard error across participants to illustrate variability only.

**Figure 14:**
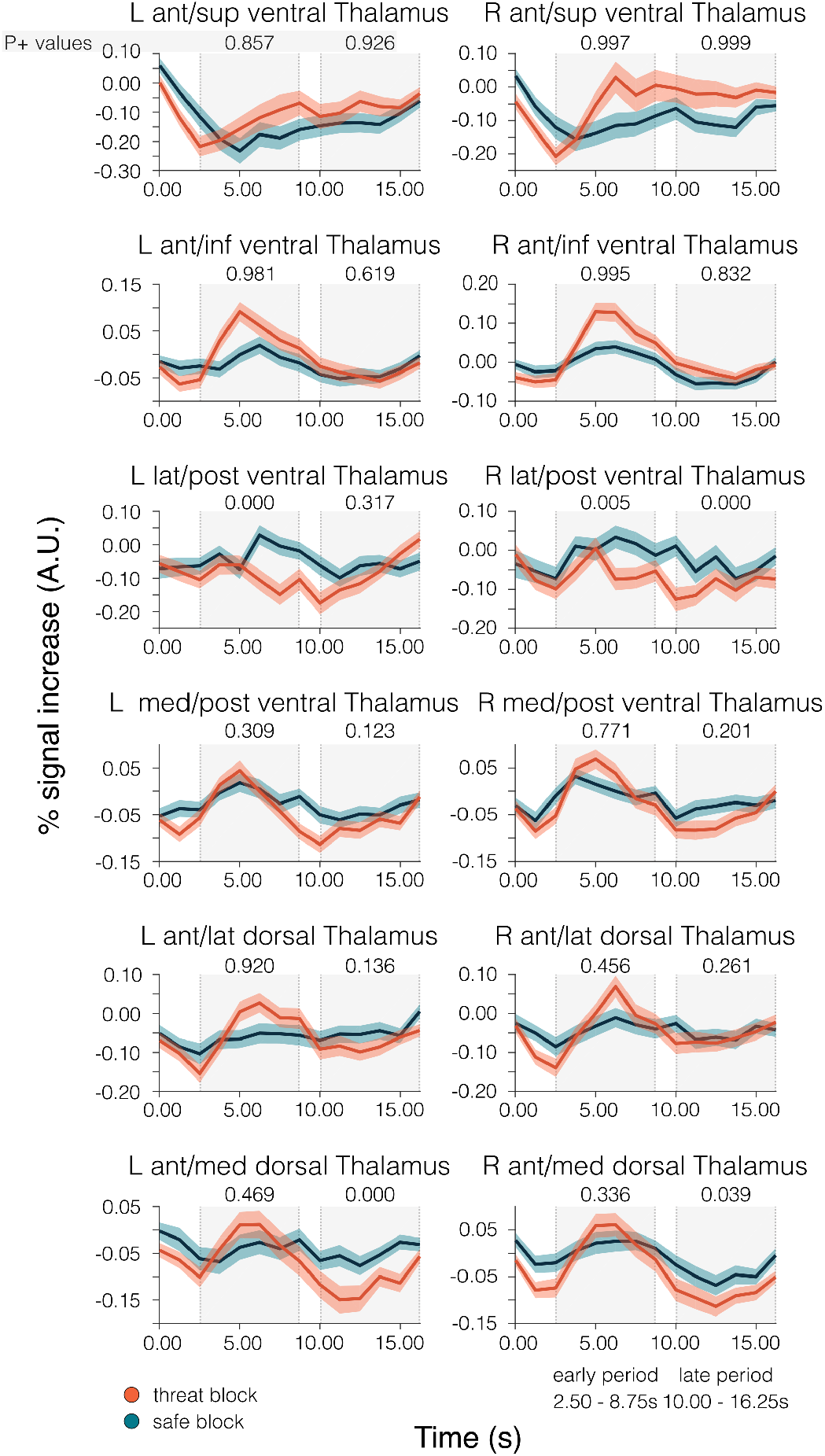
Estimated responses for thalamus. Error bands show 95% interval for standard error across participants to illustrate variability only.

### Voxel-level analysis

To characterize the spatial distribution of effects in the insula and amygdala, we performed voxelwise BML analysis. The results were in line with the ROI-level findings. For the insula (Figure 15), increased responses with `threat` were observed almost entirely in the dorsal anterior insula. In contrast, in the posterior insula, responses were stronger during `safe`. For the amygdala (Figure 16), the results followed what was found with the ROI-level analysis, but a spatial pattern of effect strength was also evident, with weaker evidence in more ventral parts of the amygdala. In other words, in the ventral amygdala, responses to `threat` and `safe` tended to be relatively similar.

**Figure 15:**
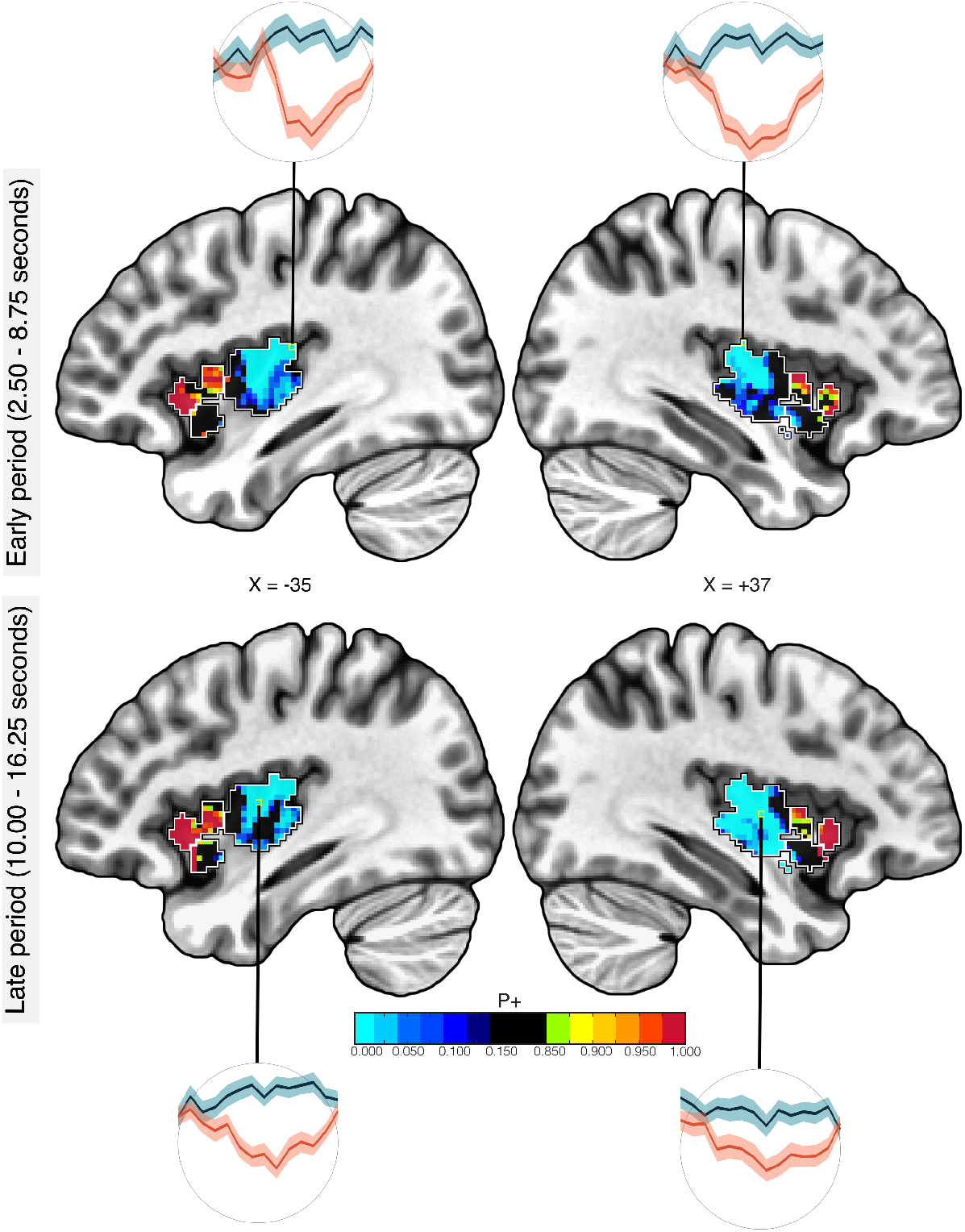
Voxelwise Bayesian multi-level model analysis: insula. Analyses were performed in the left and right insula, separately. White outlines indicate the regions employed in the ROI-level analysis. Sample <u>voxel</u> time series are shown.

**Figure 16:**
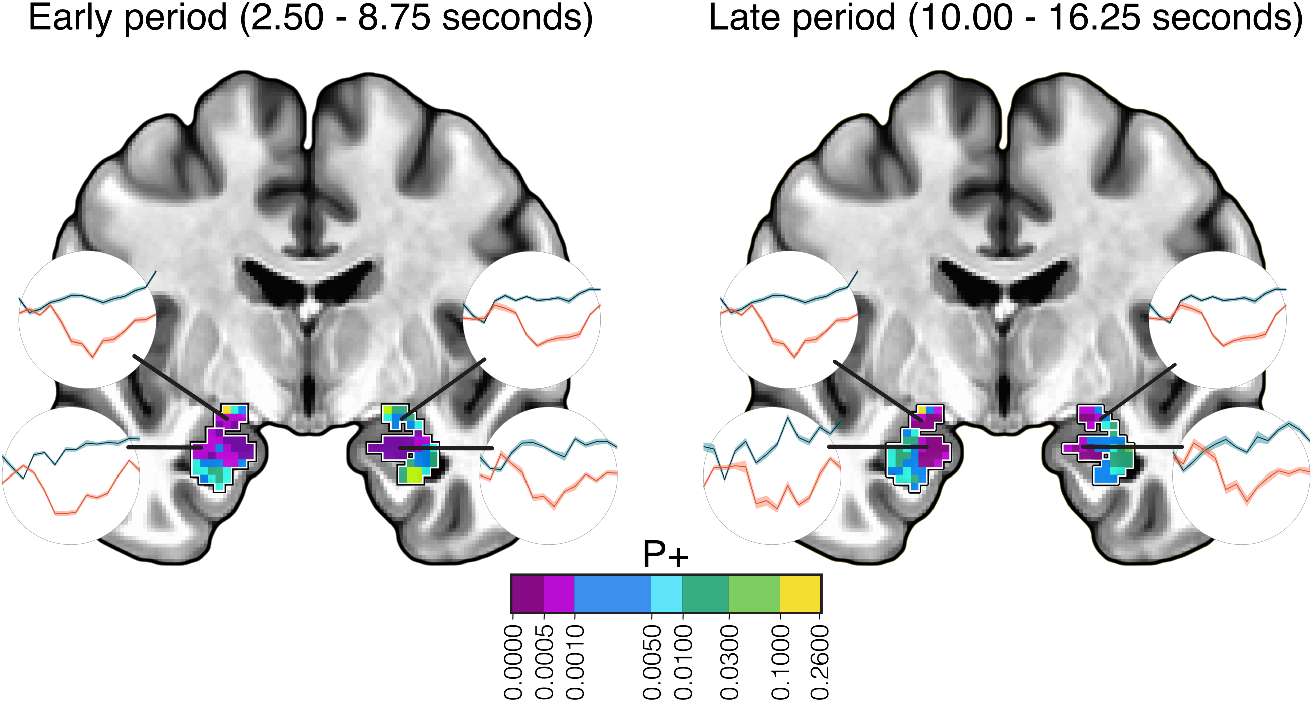
Voxelwise Bayesian multi-level model analysis: amygdala. Analyses were performed in the left and right amygdala, jointly. White outlines indicate the regions employed in the ROI-level analysis. Sample <u>voxel</u> time series are shown.

## Discussion

Sustained anticipation of unpredictable aversive events generates anticipatory processing that is central to anxiety. In the present study, we examined how sustained threat was processed in the human brain. We used a relatively large sample (*N* = 109) and employed a Bayesian multilevel analysis approach to contrast `threat` and `safe` periods. Our analyses demonstrate that the effect of sustained threat is heterogeneous and distributed across the brain. Thus, the impact of threat is widespread, and not restricted to a small set of putatively emotion-related regions, such as the amygdala and the BST. Both transient and sustained as well as increased and decreased responses were observed. See `Big Picture` box.

#### Big Picture

- Extended, uncertain threat engages the brain in a <u>distributed</u> fashion that prepares the organism to deal with the challenge. We believe that the focus on a few “typical” threat/aversive processing brain regions provides an impoverished and possibly misleading picture.
- Periods of anxious apprehension likely engage multiple mental processes, including those traditionally described as attentional, motivational, emotional, and action-related. Whereas there is value in attempting to isolate some of these processes (e.g., “purely” emotional), in more natural settings, they are <u>jointly engaged and likely intertwined</u>. We believe that always trying to disentangle them is counterproductive.
- Transitioning between `threat` and `safe` states, and vice versa, leads to a <u>massive switch in brain responding</u> likely involving most of the brain.
- Responses during `threat` and `safe` states are <u>complex and multifaceted</u>, involving both signal increase and decrease, as well as different patterns of transient and sustained signals.

##### Key findings

- Whereas some brain regions were <u>engaged</u> by threat, others were <u>disengaged</u>. Understanding both classes is necessary for a fuller appreciation of how the brain handles these states.
- Consistent with some, but not all, of the prior literature in humans, the <u>amygdala was **not** engaged</u> robustly during extended, uncertain threat.
- The <u>bed nucleus of the stria terminalis was recruited transiently</u>, not in a sustained fashion during `threat`.
- Regions that are part of the <u>default network</u> showed <u>decreased</u> responses during `threat`. It is possible that they disengage as the individual switches from self-related processing during `safe` conditions to preoccupation with the potential shock during `threat`.
- Some regions that showed decreased responses with `threat` may encode relative <u>safety</u>. Given evidence in rodents, this is possibly the case in the hippocampus.
- Responses often <u>did not conform to canonical response shapes</u>. Instead they showed nuanced/complex response profiles.

##### Caveats

- Because participants did not perform a task during `threat`/`safe` blocks, interpretation of the underlying processes is challenging.
- For example, interpretation of the responses in the posterior insula in terms of safety is speculative, despite related findings in rodents.
- For example, interpretation of the responses in the cerebellum in terms of timing mechanisms is speculative, but provide new avenues that can be investigated with experimental designs targeting this possibility.

### Threat-related increased responses

During `threat` blocks, increased responses were observed in multiple brain regions. Evidence of *transient* responses was very strong in, among others, multiple sectors of the IFG, anterior MCC, pre-SMA, dorsolateral PFC, dorsal anterior insula, thalamus, and caudate. Evidence of *sustained* responses was strongest in different sectors of the IFG, dorsal anterior insula, posterior MCC, dorsolateral PFC, and putamen.

An extensive literature has described the engagement of the <u>anterior MCC</u> in aversive processing (for a review see, (Vogt, 2005)). ^2^ Current evidence supports the idea that the anterior MCC is involved in emotion-, cognition-, and pain-related processing (Shackman et al., 2011); see also, (Misra and Coombes, 2015). In addition, meta-analyses suggest that the anterior MCC plays a central, integrative role in emotion regulation (Kohn et al., 2014), and is part of a core system for implementing self-control across emotion and action domains (Langner et al., 2018). The present study adds to this literature by showing that this region is transiently engaged when a threat context is initially experienced, and closely replicates a previous finding by our lab ((McMenamin et al., 2014); see also (Hasler et al., 2007; Schlund et al., 2013)). ^3^

The dorsal <u>anterior insula</u>, particularly on the right, exhibited sustained responses that were greater for `threat`, consistent with other empirical findings (Alvarez et al., 2011; Somerville et al., 2013). Indeed, several conceptual frameworks implicate the anterior insula in the processing of sustained threat, and have suggested that impaired processing of this area underlies anxiety disorders (Paulus and Stein, 2006; Picó-Pérez et al., 2017). Notably, we observed enhanced responses during `threat` only in the <u>dorsal, not ventral</u> anterior insula. These results are in contrast with proposals that suggest that the ventral anterior insula is more tuned to emotion-related processing whereas the dorsal sector is more tuned to cognitive-related processing (Kurth et al., 2010; Uddin et al., 2017).

In the present study, the <u>dorsolateral PFC</u> showed sustained activation during threat. Although this brain sector is often associated with cognitive functions, it is also involved during affective processing. Consistent with our results, studies have reported sustained signals lasting up to 30 seconds in the dorsolateral PFC (Hur et al., 2020; Andreatta et al., 2015; Wood et al., 2015; Qin et al., 2009). ^4^

Surprisingly, responses in the <u>BST</u> were relatively transient, whereas we had expected them to exhibit sustained responses during the block period given evidence of more sustained responses in this region (Alvarez et al., 2011; Somerville et al., 2013; McMenamin et al., 2014; Hur et al., 2020). Other regions exhibited sustained responses, including the right dorsolateral PFC, the right anterior putamen, among others. ^5^

Although threat-related research tends to focus on a few select structures (e.g., amygdala and medial PFC), there are increasing efforts to understand how distributed/large-scale circuits are involved. For example, in a review of fear and anxiety, Lüthi and colleagues (Tovote et al., 2015) described the participation of the amygdala, hippocampus, hypothalamus, PAG, thalamus, BST, medial PFC, and ventral striatum (including subregions of these structures), among others. Furthermore, it has been proposed that circuits involving multiple amygdala nuclei, PAG, medial PFC, and ventral striatum/nucleus accumbens participate in resolving between escape and freezing behavioral alternatives (LeDoux et al., 2017; Moscarello and LeDoux, 2013; Moscarello and Hartley, 2017); see also (Ilango et al., 2014).

In this context, it is noteworthy that we observed robust responses to `threat` in the <u>striatum</u>, in both the anterior caudate and the anterior putamen. The involvement of the anterior putamen was sustained, and to a some extent that of the anterior caudate, too. Note that because the block period did not require any response, and because both conditions required the same final type of motor response, it is unlikely that the differential response was substantially driven by motor-related processing. In this regard, the distinct response pattern in the posterior caudate is relevant, as it exhibits a tendency to be stronger for `threat` only during the late period.

Our results revealed a diverse set of response patterns across nuclei of the <u>thalamus</u>. Both anterior nuclei (superior and inferior) of the *ventral thalamus* exhibited stronger responses for `threat`, which were particularly robust and sustained in the right hemisphere. This pattern contrasted to that of the medial and lateral nuclei of the ventral thalamus.

### Threat-related decreased responses

Multiple brain regions respond vigorously when threats are encountered or experienced. But it is also noteworthy that multiple brain regions respond <u>less strongly</u> during threat relative to safe, as observed here and in other studies (Limbachia et al., 2021; Mobbs et al., 2010). In our study, because no task was required during blocks, it is possible that decreased responses were related to routine processing in <u>default network</u> regions. The idea is that a region (say, PCC) is engaged during periods without an overt task, such as during safe blocks. However, during a threat block, these regions are not recruited as much, as the individual is now preoccupied with the potential shock.

It is noteworthy that some of the regions exhibiting decreased signals did not overlap with the default network. ^6^ For example, our medial OFC ROI was entirely more ventral relative to parts of the medial OFC that are observed in the default network. The ROI that we called `IFG2` (see Figure 2) had very minimal overlap (2-3 voxels) with the default network. In addition, the posterior insula showed marked decreases. Among subcortical regions, decreased responses were observed in the ventral thalamus, posterior caudate, and parts of the cerebellum (see below) not typically linked to the default network. Combined, these results indicate that several brain regions are less engaged during `threat` relative to `safe` not because they are part of the default network but due to other aspects potentially linked to safety-related processing (see below for further discussion of the posterior insula).

The status of amygdala responses to <u>sustained threat</u> in humans is unclear. Whereas some investigators have reported increased responses (Schlund et al., 2013), several others have actually observed decreases (Choi et al., 2012; McMenamin et al., 2014; Pruessner et al., 2008; Wager et al., 2009). Here, we observed weaker responses for `threat` in all amygdala ROIs. Furthermore, the response evolution indicated that signals during `threat` decreased relative to `safe` during the early period, and remained lower during the late period. The voxelwise analysis confirmed these results, demonstrating that they did not result from the ROI-averaging process. At the same time, the analysis showed that the most robust differences were observed in relatively dorsal parts. Thus, in line with several other imaging studies, under the conditions of extended, uncertain threat of our experiment, the amygdala did not exhibit increased responses (but see Hur et al. 2020); instead the central/medial amygdala and basal amygdala exhibited a slowly evolving decreasing response. In passing, we not that while the amygdala is routinely associated with threat and fear-related processing, studies have observed safety-related signals in the basal amygdala using classical conditioning paradigms in both rats (Sangha et al., 2013) and non-human primates (Genud-Gabai et al., 2013).

In a previous study of threat controllability in humans, we observed responses in the <u>posterior insula</u> that were stronger during controllable relative to uncontrollable shocks (Limbachia et al., 2021). These findings were notable because in a threat controllability study in rodents, the posterior insula was suggested to encode a `safety signal` (Christianson et al., 2011). In the present study, the voxelwise Bayesian analysis identified voxels in the posterior insula with responses greater for `safe` relative to threat (also seen in the ROI-based analysis). The temporal evolution of the responses was also quite noteworthy, as the responses increased early on for both `threat` and `safe`, rapidly decreased for `threat` but stayed elevated for `safe` until the end of the block (the same pattern was observed in the ROI-based analysis). Combined, these results are consistent with the notion that the posterior insula potentially signals a state of relative safety.

Research over the past few decades has revealed that the <u>cerebellum</u> contributes to an array of non-motor functions, even including fear conditioning (Fullana et al., 2018). Lobule VI ROIs showed transient responses during `threat` blocks with rather modest increases relative to `safe` blocks (Figure 17; see also (Hur et al., 2020)). A qualitatively different pattern of results was observed in the Crus I ROIs, where responses were <u>weaker</u> for `threat` during the early period. Intriguingly, the responses appear to evolve with a somewhat ramping activity. It is noteworthy that a similar area of the cerebellum was reported to be involved in time perception and anticipation of future events (Apaydın et al., 2018). More broadly, the cerebellum is implicated in motor timing and planning (see Tanaka et al. 2020 for review). We speculate that the responses could be related to the anticipation of the anxiety rating following the block. Finally, we note that the Crus I ROIs overlaps with sites that are correlated with the executive network during task-free conditions (Habas et al., 2009), highlighting the importance of further investigating cerebellar contributions to diverse mental processes.

**Figure 17:**
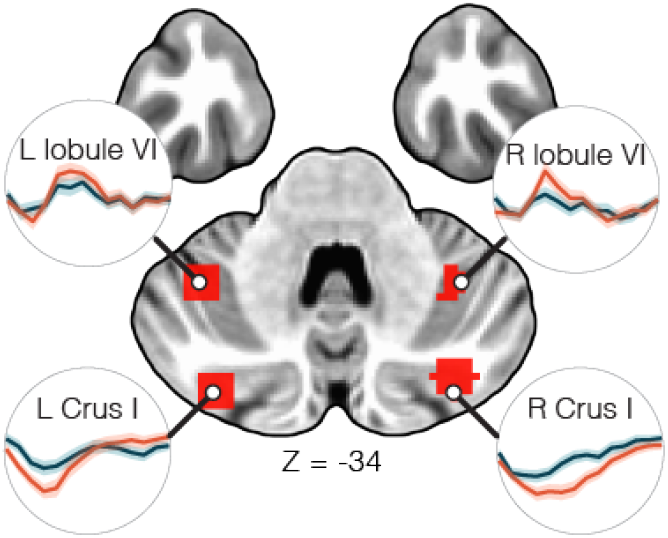
Cerebellum responses. Horizontal slice through the base of the brain showing the location of cerebellar ROIs, and the estimated ROI responses averaged across voxels during `threat` (red) and `safe` (blue). Error bands show 95% interval for standard error across participants to illustrate variability only.

### Conclusions

The present study investigated sustained threat-related processing across brain regions previously discussed in the threat-related literature. We employed Bayesian multilevel modeling to estimate effect strength when comparing `threat` versus `safe` periods. Our study showed that periods of uncertain threat engage a broad, distributed set of brain regions, which displayed a diverse set of response profiles, including transient and sustained, increased or decreased responses. Periods of anxious apprehension are likely to engage a number of mental processes, including those traditionally described as attentional, motivational, emotional, and action-related, among others (but see (Cisek, 2019; Pessoa et al., 2021). Whereas there is value in attempting to isolate some of these processes (say, the more “purely” emotional), in more natural settings, they are jointly present and likely intertwined. In this context, our experiment uncovered a rich repertoire of response patterns during both `threat` and `safe` states, showing the multifaceted ways in which the brain is engaged during these two states.

## Acknowledgements

We acknowledge Chirag Limbachia for work on data analysis during early stages of this project, and thank Gang Chen for discussions concerning statistical analysis. Research was supported by the National Institute of Mental Health (R01 MH071589).

## Footnotes

1 Readers may choose to apply thresh-olds, although we find such approach unproductive; For in-depth discussion, see (Chen et al., 2020, 2021).

2 In sharp contrast with the notion that the rostral ACC and the anterior MCC (also called dorsal ACC in some studies) are specialized in terms of emotional and cognitive processes, respectively (Bush et al., 2000).

3 But see (Alvarez et al., 2011) and (Hur et al., 2020) for evidence of a more sustained response profile.

4 In our experiment, the argument could be made that the involvement of the dorsolateral PFC is more related to “sustained attention” than threat-related processes. This is a valid point, but see `Big Picture` for further discussion.

5 Note that the skin conductance level was not sustained throughout the block duration either (see Figure 6). However, transient skin conductance responses were not due to uneven distribution of shocks, as these events were equally distributed throughout the block, including towards the end of the block.

6 To probe this issue for cortical regions, we investigated potential overlap by comparing our ROIs to the masks available in the so-called Schaefer parcellation (Schaefer et al., 2018).

